# Decoding the fibroblast/mast cell signaling pathway of acupuncture

**DOI:** 10.1101/2025.11.20.688774

**Authors:** Xiqing Xue, Rui Wang, Wenru Sheng, Jie Pang, Dongyu Lu, Meng Yu, Tian Lan, Yiider Tseng

**Affiliations:** The Innovation Institute of Chinese Medicine, Shandong University of Traditional Chinese Medicine, Jinan, Shandong, 250355, China; Institute of Acupuncture and Moxibustion, Shandong University of Traditional Chinese Medicine, Jinan, 250355, China; College of Traditional Chinese Medicine, Shandong University of Traditional Chinese Medicine, Jinan, Shandong, 250355, China

## Abstract

Acupuncture has been practiced for thousands of years with documented therapeutic effects, yet its underlying biological mechanisms remain poorly understood. Here, we show that acupuncture stimulation converts local mechanical forces into neuronal signals through defined cellular interactions, establishing a causal axis that links micro-level events at the ST36 acupoint to systemic therapeutic effects. We demonstrate that acupuncture needle manipulation transmits mechanical tension through collagen fibers to local fibroblasts, which subsequently secrete stem cell factor (SCF) and interleukin-33 (IL-33) in the acupoint. SCF recruits mast cells, while IL-33 induces their activation and the release of neuromodulatory molecules. These molecules then engage the nervous system, as evidenced by c-fos expression in the lumbar dorsal horn, demonstrating the transmission of acupuncture-induced peripheral signals into central circuits. Together, these findings define a functional axis linking needle-evoked stimulation to neuronal activation and provide a mechanistic framework for understanding the biological basis of acupuncture and for its evidence-based refinement.

---

Acupuncture has been a cornerstone of traditional Chinese medicine for more than two millennia and is practiced worldwide for a variety of conditions. Global attention to acupuncture rose in the 1970s after reports of its use as an alternative to surgical anesthesia, and archaeological discoveries such as the 5300-year-old Otzi the “Iceman” in the Italian Alps, whose skin contained holes corresponding to traditionally documented acupoints (*i.e.*, the points on the skin for acupuncture practice to take effects)^1–3^, highlight the practice’s deep historical originals. The presence of these separate acupoints through time and space raises the question of the benefits of acupuncture, and whether its therapeutic outcomes can be explained by defined biological mechanisms. Clinical studies have reported therapeutic benefits of acupuncture across a range of conditions^4–11^. However, the biological mechanisms of needle insertion and manipulation at acupoints remain poorly understood.

Acupuncture is generally categorized into two forms of stimulation: traditional manual acupuncture and electroacupuncture. The latter delivers electrical pulses to acupoints through the needle, directly exciting local neurons through deep connective tissue^12,13^. Recent research has shown that activation of PROKR2^Cre^-labeled neurons in the myofascia at the ST36 (Zusanli) acupoint can trigger vagus nerve discharge and suppress lipopolysaccharide (LPS)-induced inflammation, establishing foundational scientific evidence for the principles of electroacupuncture^14,15^. Traditional acupuncture, by contrast, engages subtle mechanical forces generated by needle insertion to acupoints, which elicit therapeutic behaviors^16,17^. These mechanical inputs represent a distinct route for engaging cellular and neuronal responses, uncovering the paths through which acupuncture stimuli give rise to integrated physiological effects.

These pathways also provide greater insight into the routes through which acupuncture elicits its initial response at each acupoint. Understanding the process helps clarify how the signal propagates and integrates within the nervous system, followed by modulation of these signals in target tissues and organs. In an effort to further elucidate this process and complement the significant advances already made in understanding the interactions between the brain and peripheral organs and tissues^18–27^, we focus here on the initial stages to explore the working principles and mechanisms of acupuncture at the acupoint level.

During traditional acupuncture (henceforth, “acupuncture”), manual insertion of a needle into an acupoint generates local mechanical forces that ultimately elicit therapeutic responses through the nervous system^28–30^. Recent evidence suggests that mast cells contribute critically to the analgesic effects observed with acupuncture stimulation^31–34^. In a rat model, mast cell degranulation at the ST36 acupoint is associated with enhanced discharge of the sciatic nerve and subsequent analgesia^35^. Conversely, administration of the mast cell stabilizer sodium cromoglicate (DSCG) suppresses mast cell degranulation, normalizes sciatic nerve activity, and markedly attenuates the analgesic response. These findings implicate mast cell activation as a key mediator in the conversion of local acupuncture stimulation into neural signals that underlie analgesic outcomes.

While these findings establish a critical role for mast cells in acupuncture, defining how mechanical stimulation from needle insertion activates mast cells and shapes the therapeutic effects of specific acupoints is crucial for translating acupuncture into targeted therapies for conditions, such as inflammation and anesthesia. Here, we address this question by uncovering the mechanisms operating at the ST36 acupoint.

## Fibroblasts are crucial for acupuncture

Recent literature has elucidated the mechanisms through which electrical stimulation in electroacupuncture evokes neuronal excitation at the ST36 acupoint. However, the analogous mechanism in traditional acupuncture remains a mystery (**Fig. 1a**). We first examined the effects of acupuncture needle insertion using ultrasound (**Extended Data Fig. 1**). The images revealed that needle manipulation induces direct mechanical interactions with collagen fibers. Fibroblasts both produce collagen and are physically anchored to the collagen matrix^36,37^, suggesting that they may be among the first cells at acupoints to detect the mechanical forces generated by acupuncture.

**Figure 1.**
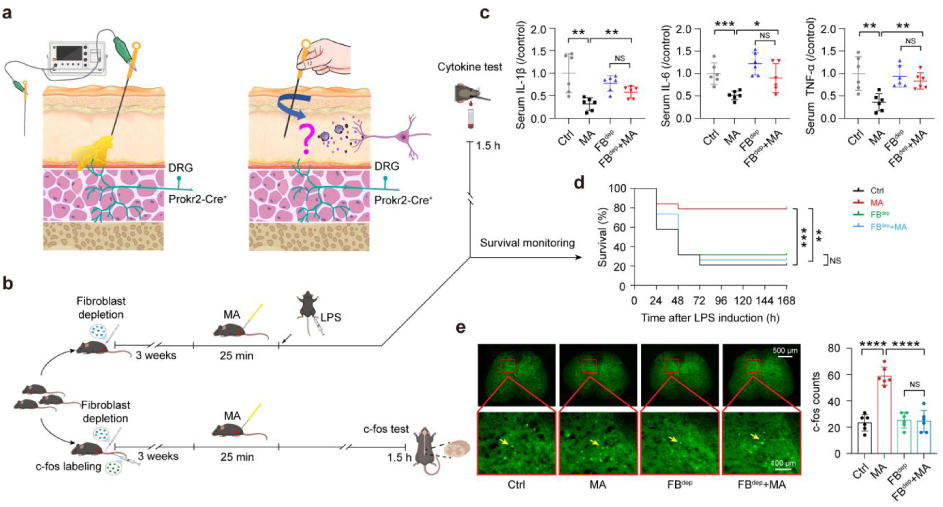
Fibroblasts are essential in the ST36 acupoint for manual acupuncture to take effect in preventing LPS-induced endotoxemia in mouse model. a. Schematic of the ST36 acupoint under electrical (left) or manual (right) acupuncture. b. Schematic of the experimental procedure. c. IL-1β/IL-6/TNF-α level in mice serum 90 minutes after LPS induction (n = 6). Ctrl: only performed LPS induction; MA: manual acupuncture; FB^dep^: fibroblast depletion; FB^dep^+MA: fibroblast depletion with MA. d. The survival rate of mice after LPS induction, recorded for 168 hours (n = 20). Significance was determined using Mantel-Cox log-rank test. e. Images (left) and data analysis (right) of c-fos expression (green) in the dorsal horn of the lumbar spinal cords (n = 6). Scale bars: 100 μm. Unless otherwise specified, data in all figures are geometric mean ± s.d.; all significance were determined using two-side Student’s unpaired t-test; *, **, ***, and **** denote *p* < 0.05, 0.01, 0.001, and 0.0001, respectively. NS denotes non-significant, *p* > 0.05.

Thus, we depleted fibroblasts in the ST36 acupoint to examine the role of fibroblasts in preventing LPS-induced inflammation (endotoxemia) in mice. We examined the levels of the represented inflammatory cytokines in the serum and the survival rate in each of the cases. We randomly divided 128 mice into four groups: Control (Ctrl); manual acupuncture (MA); fibroblast depletion (FB^dep^); and fibroblast depletion with MA (FB^dep^+MA) (**Fig. 1b**). The mice in the two groups marked FB^dep^ received AAV injections at the ST36 acupoint to deplete fibroblasts (**Extended Data Fig. 2**), while the other two groups received equivalent injections of phosphate-buffered saline (PBS). We further randomly selected six mice from each group, which were subjected to an intravenous injection of an AAV to fluorescently label the new expression of c-fos in the lumbar dorsal horns, a marker of neuronal activation (**Extended Data Fig. 3**).

Three weeks later, mice in the MA and FB^dep^+MA groups received manual acupuncture at the ST36 acupoint, while Ctrl and FB^dep^ groups remained untreated. Ninety minutes after stimulation, lumbar spinal cords were collected from the six mice per group that had received intravenous injection to assess the c-fos expression in the dorsal horn (n = 6). In parallel, mice not used for c-fos analysis received an intraperitoneal LPS injection 25 minutes after acupuncture to induce endotoxemia. Another 90 minutes later, whole-blood samples from six mice per group were collected to measure inflammatory cytokine levels (n = 6), and the remaining mice were monitored to determine survival rates (n = 20).

As the serum test results were obtained immediately after LPS induction, we began by assessing the serum concentrations of the inflammatory cytokines IL-1β, IL-6 and TNF-α in the four groups of mice following LPS-induced endotoxemia (**Fig. 1c**). By examining the serum cytokine levels, we found that these were significantly lower in the MA group than in the Ctrl group (at least *p* < 0.01), supporting the anti-inflammatory efficacy of acupuncture. In addition, the serum cytokine levels in the FB^dep^+MA group were not significantly different from those in the FB^dep^ group and remained significantly higher than in the MA group (at least *p* < 0.05), demonstrating that fibroblasts play a critical role in mediating acupuncture’s suppression of systemic inflammation. Fibroblasts at the acupoint are therefore essential for the therapeutic efficacy of acupuncture in endotoxemia. Furthermore, these results also support that acupuncture can attenuate LPS-induced inflammation in mice and validate the LPS-induced endotoxemia model as a suitable platform for mechanistic studies of acupuncture.

One week later, we obtained the survival rate results of mice across the different groups (**Fig. 1d**). The survival rate of mice in the MA group was significantly higher than that in the Ctrl group (*p* < 0.001), demonstrating the protective effect of acupuncture. By isolating the effects of fibroblasts, we find that the removal of fibroblasts eliminated the benefit of acupuncture: survival in the FB^dep^+MA group did not differ significantly from the FB^dep^ group, and the FB^dep^+MA group was significantly lower than the MA group (*p* < 0.01).

Ultimately, for acupuncture to be therapeutically effective, signals initiated at acupoints must be transmitted to the nervous system^37^. To test whether fibroblast depletion disrupts this process, we examined lumbar spinal cords for c-fos expression following acupuncture at the ST36 acupoint (**Fig. 1e**). Robust c-fos induction in the dorsal horn was observed only in mice with intact fibroblasts, indicating that fibroblasts are required for transmission of acupuncture signals to the dorsal horn.

Taken together, these findings demonstrate that fibroblasts at the acupoint are indispensable for acupuncture, as they are required for its protective effects against endotoxemia, suppression of systemic inflammation, and activation of spinal neurons.

## Fibroblasts facilitate acupuncture

Although mast cells are widely regarded as an intermediary for transmitting the acupuncture signals from the acupoints to the nervous system, it remains unclear whether they serve as the primary responders to the force stimuli generated from acupuncture. To investigate this relationship, we examined the responses of mast cells to mechanical stimuli analogous to those elicited from acupuncture.

Acupuncture transmits forces along collagen fibers, generating local tension; to replicate an analogous environment, we adapted a stretching device to mimic the mechanical stimuli present at the acupoint during acupuncture. To isolate the effects of the mast cells, we load mast cells onto the surface of the device coated with collagen. After one day of incubation, we applied stretching motion (5% or 10% of amplitude; 0, 0.5 or 2 Hz of frequency) to the device for 30 minutes to mimic the acupuncture process (**Fig. 2a**). Once complete, the supernatant of the culture media was collected from the stretching device and subjected to the enzyme-linked immunosorbent assay (ELISA) to examine the release of the neuromodulatory molecules, such as neuropeptides (represented by substance P (SP)) and neurotransmitters (represented by 5-Hydroxytryptamine (5-HT), also called serotonin), from mast cells. The results indicate that mast cells were unresponsive to the range of stretching forces, suggesting that the mechanical stimuli generated by the acupuncture procedure is not the direct trigger for mast cells to release neuromodulatory molecules. This implies that other cells types are involved in converting mechanical stimuli into signals that mast cells can respond to.

**Figure 2.**
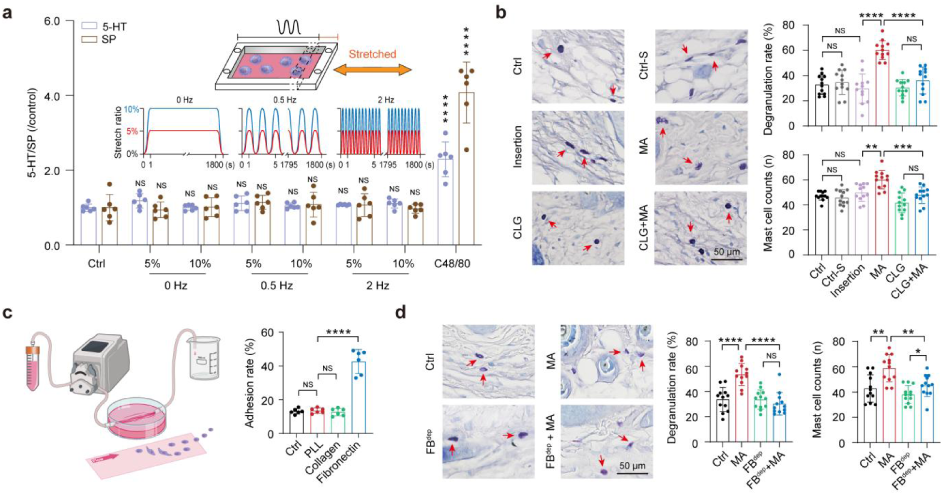
Fibroblasts play a critical role in releasing neuromodulatory factors on mast cells during acupuncture. **a.** Schematic of a stretching device. The device can be controlled to provide the simple-harmonic-motion stretches. Substance P (SP) and 5-hydroxy-tryptamine (5-HT) levels measured in cell-free, supernatants of P815 mast cells after being stretched for 30 minutes (amplitude: 5% or 10% ; frequency: 0, 0.5 or 2 Hz) (n = 6). **b.** Toluidine blue staining on the ST36 acupoint tissue section. Ctrl: Normal condition; Ctrl-S: saline injection; Insertion: needle only inserting into the acupoint, no rotation; MA: manual acupuncture (insertion and rotation); CLG: collagenase injection; CLG+MA: collagenase injection with MA. Scale bar: 50 μm. The number and the degranulation rate of mast cells were analyzed (n = 12). **c.** Schematic of microfluidic device detecting mast cell adhesion ability (left). The dishes were left blank (Ctrl), coated by poly-L-lysine (PLL), collagen, or fibronectin. Statistical chart of the adhesion ability of mast cells with different substrates (right). **d.** Toluidine blue staining on the ST36 acupoint section (left). Ctrl: without manipulation; MA: manual acupuncture; FB^dep^: fibroblast depletion; FB^dep^+MA: fibroblast depletion with MA. Scale bar: 50 μm. Statistics of the level and degranulation rate of mast cells in the ST36 acupoint (n = 12) (right). Please note, for b. and d., as each mouse has two ST36 acupoints, the number of the acupoint samples is twice the mice number.

The ultrasonic videos reveal that collagen fibers are the main structures in contact with the acupuncture needle; hence, we injected collagenase into the ST36 acupoint to digest collagen fibers before acupuncture and examined whether the quantity and degranulation rate of mast cells still increases during acupuncture (**Fig. 2b**). Comparing the biopsied tissue sections (dyed with toluidine blue) under the normal condition *vs.* the collagenase treatment condition reveals that collagen digestion prevents the quantity of mast cells from increasing and degranulating. This suggests that the ST36 acupoint requires mechanical force propagation through collagen fibers for mast cells to respond to the acupuncture.

Having established the essential role of fibroblasts in acupuncture, we proceeded to observe the distribution and morphological alterations of both fibroblasts and mast cells by immunostaining tissue sections from the ST36 acupoint of mice before and after acupuncture using antibodies against collagen, vimentin (a marker of fibroblasts and mast cells) and c-kit (a marker of mast cells) (**Extended Data Fig. 4**). After acupuncture, both the number of mast cells and the rate of degranulation appeared to increase in the ST36 acupoint. While it is difficult to determine whether mast cell (co-labeled by vimentin and c-kit) and the nearest fibroblast (labeled by vimentin) engaged with each other, we can conclude that fibroblasts and mast cells either interact or remain close after acupuncture.

To evaluate whether the mast cells physically interact with fibroblasts at the acupoint during acupuncture stimulation, we used a microfluidic device to generate a shear flow to carry mast cell cultures forward; in front, the culture surface was left blank, coated with poly-L-lysine, collagen, or fibronectin to examine the attachment capacity of mast cells (**Fig. 2c**). As mast cells adhere specifically to fibronectin surfaces, we performed fluorescence immunostaining for fibronectin in tissue sections and fibroblast cultures to examine whether fibronectin mediates fibroblast-mast cell contact (**Extended Data Fig. 5**). The images suggest that settled fibroblasts are fully covered by fibronectin in both conditions; hence, the mechanical stimulation of acupuncture likely promotes fibroblast-mast cell contact through fibronectin-mediated adhesion.

To further evaluate whether mast cell chemotaxis and degranulation are related to fibroblasts, we examined the acupuncture-induced mast cell activities under fibroblast depletion in the ST36 acupoint. The mice were randomly separated into four groups: Ctrl, MA, FB^dep^ and FB^dep^+MA (n = 6). Mice in the FB^dep^ and FB^dep^+MA groups were injected the specific AAVs at the ST36 acupoint to induce fibroblast apoptosis, while the other two groups were injected with PBS. Three weeks later, the MA and FB^dep^+MA groups were treated with acupuncture. After 90 minutes, all groups were biopsied at the ST36 acupoint with toluidine blue staining (**Fig. 2d**). Micrographs revealed that acupuncture increased mast cell numbers only when fibroblasts were present in the acupoint. In addition, mast cell degranulation was also significantly elevated in the presence of fibroblasts.

Taken together with the essential role of fibroblasts in acupuncture function, our results suggest that acupuncture-induced mechanical stimuli most directly impact the fibroblasts, which then serve as an intermediary channel through which mast cells trigger chemotaxis and degranulation.

## Fibroblasts convert mechanical signals

To further detail the role of fibroblasts in mediating mast cell activity during acupuncture, we examined their response under force stimuli.

Previous reports have shown that fibroblasts in connective tissue become broader and thicker as a result of acupuncture^17,38^. We therefore used this cell morphological change as a baseline to determine the degree of stretching *in vitro*. Fibroblasts cultured on the fibronectin-coated stretching device were subjected to different amplitudes of stretching (**Extended Data Fig. 6**). A 5% stretch produced similar morphology changes in fibroblasts compared to those in the connective tissue explant. A 10% stretch caused partial detachment of cells from the surface.

Hence, we used 5% stretching to evaluate the fibroblast-mediated effects on mast cells. The supernatant from the stretched fibroblast culture (SSFC) was applied to the mast cell culture, and the mast cell supernatant was subsequently subjected to ELISA against SP and 5-HT (**Fig. 3a**). Mast cells exposed to SSFC released significantly higher levels of SP and 5-HT, indicating that fibroblast-derived factors stimulate mast cell activation.

**Figure 3.**
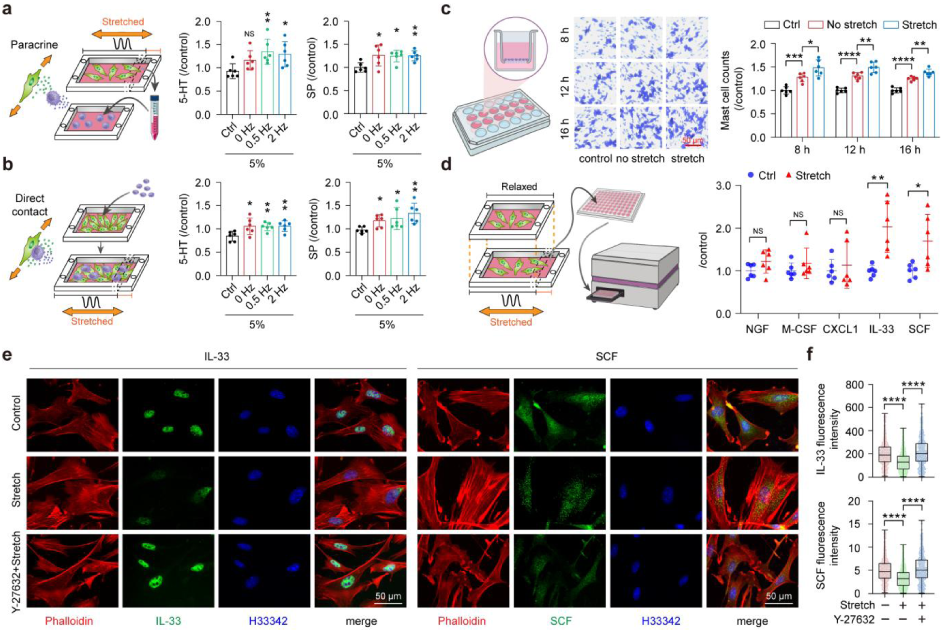
Fibroblasts orchestrate mechanotransduction of acupuncture signals at acupoints. **a.** Schematic of stretched fibroblasts on mast cells through paracrine signaling. Substance P (SP) and 5-hydroxytryptamine (5-HT) released by P815 mast cells were measured after culturing in the supernatant of stretched fibroblasts (5%; 0, 0.5 or 2 Hz; 30 minutes) for 4 hours (n = 6). **b.** Schematic of mechanical force on P815 cells, co-cultured with fibroblasts. SP and 5-HT levels in the co-culture supernatant after stretching (5%; 0, 0.5 or 2 Hz; 30 minutes) (n = 6). **c.** Schematic of the Transwell device (left): P815 cells were seeded in the upper chamber, with the lower chamber containing complete medium (Ctrl), conditioned medium from non-stretched fibroblasts (No stretch), or conditioned medium from stretched fibroblasts (Stretch). Images of cell clustering were stained with crystal violet after 8, 12 or 16 hours of incubation (middle). Scale bar: 50 μm. Statistical chart of mast cells under chemotaxis (n = 6) (right). **d.** Schematic of cytokine levels in the supernatant of fibroblasts after stretching by ELISA (left). NGF/CXCL1/M-CSF/IL-33/SCF levels measured in cell-free supernatants of fibroblasts after stretching (5%; 2 Hz) for 30 minutes (n = 6). **e.** Immunofluorescence staining of mice skin primary fibroblasts, with and without Y-27632, pre- and post-stretch: phalloidin (red), nucleus (blue), IL-33 or SCF (green). **f.** Fluorescence intensity statistics of IL-33 (top) and SCF (down) in fibroblasts. For IL-33, Ctrl (n = 507), Stretch (n = 493), Y-27632 with Stretch (n = 452). For SCF, Ctrl (n = 255), Stretch (n = 359), Y-27632 with Stretch (n = 404). Scale bars are 50 μm.

Since cell contact may contribute to mechanical stimulation, fibroblasts and mast cells were further co-cultured for 4 hours to permit direct contact, and the co-culture was subjected to 5% stretching for 30 minutes (**Fig. 3b**). ELISA of the supernatant revealed that co-culture under stretching increased SP and 5-HT release, compared to that without stretching. Therefore, it is also possible that acupuncture-induced tissue deformation promotes interactions between fibroblasts and mast cells at the acupoint, thereby amplifying the release of neuromodulatory molecules from mast cells.

To understand the cause of mast cell chemotaxis at acupoints following acupuncture, we next tested the chemotactic capacity of SSFC to attract mast cells. We measured mast cell chemotactic levels using a Transwell device, where a thin, porous layer separates the upper and lower chamber. The mast cells were placed in the upper chamber and SSFC was added to the lower chamber (**Fig. 3c**). SSFC significantly increased mast cells passing through the porous layer relative to supernatants from the unstretched fibroblast culture, indicating that SSFC contains chemotactic factors. This finding provides an explanation for the accumulation of mast cells at acupoints following acupuncture.

These results consistently indicate that the mechanical signals introduced by acupuncture first stimulate fibroblasts, which transmit signals downstream to mast cells in the acupoints. Therefore, we explored the candidate molecules released from stretched fibroblasts targeting mast cells through comparing transcriptomic sequencing between fibroblasts under normal and stretching (5% amplitude and 1-Hz frequency for 30 minutes) conditions (n = 3) (**Extended Data Fig. 7**). Accordingly, we identified 1567 differentially expressed genes whose expression levels differed significantly after cells were subjected to stretching (*p* < 0.05). In addition, most of these detected genes are up-regulated following stretching.

We next performed Gene Ontology (GO) and Kyoto Encyclopedia of Genes and Genomes (KEGG) pathway enrichment analyses through the STRING database. The results suggest that during the thirty minutes of mechanical stretching, fibroblasts’ responses in protein expression remain focused on the ribosome biogenesis and protein translation stages, rather than progressing towards the synthesis of new secretory proteins.

These findings imply that under the present conditions, acupuncture-like mechanical stimulation may primarily facilitate the release of pre-existing proteins, rather than induce *de novo* synthesis of secretory factors. Accordingly, the effective molecules from fibroblasts driving mast cell chemotaxis and releasing neuromodulatory molecules, such as SP and 5-HT, must already exist inside the fibroblasts. To our best knowledge, fibroblasts can secrete neuron growth factor (NGF), chemokine (C-X-C motif) ligand 1 (CXCL1; also called growth-regulated oncogene α; GROα), macrophage colony-stimulating factor (M-CSF), interleuking-33 (IL-33) and stem cell factor (SCF) to induce mast cells releasing neuromodulatory molecules under certain circumstances. To test the mechanical stimuli on fibroblast secretion, the SSFCs were collected and analyzed by ELISA for those molecules (**Fig. 3d**). The results showed that only IL-33 and SCF concentrations changed significantly in response to stretching relative to unstretched cultures.

To examine how mechanical stretching alters the distributions of IL-33 and SCF in the fibroblasts, we performed immunofluorescence staining on these two molecules in fibroblasts before stretching, after stretching, and after stretching under the ROCK inhibitor Y-27632, which reduces cellular tension (**Fig. 3e**).

The micrographs showed that IL-33 was originally localized predominantly inside the nucleus and perinuclear region of the fibroblasts, but stretching caused a marked decrease in nuclear and perinuclear intensity, indicating secretion into the extracellular space. In addition, the presence of Y-27632 can effectively prevent the stretching influence to the cells. In the case of SCF, stretching transformed the original spreading distribution of SCF in the cells into sparkling granules, and the overall SCF intensity in the cells decreases significantly, compared to the un-stretched cells and the cells subjected to Y-27632.

We compared the intensities of IL-33 and SCF inside the cells before stretching, after stretching, and after stretching under the presence of Y-27632 (**Fig. 3f**). The results clearly demonstrated that the IL-33 and SCF signaling diminish from the fibroblasts after acupuncture, indicating their secretion out of the fibroblasts.

## SCF and IL-33 in acupuncture

Given that SCF and IL-33 stood out as potential regulators of acupuncture, we next examined whether either molecule alone could induce mast cells to release neuromodulatory molecules. SCF and IL-33 were separately added into mast cell cultures; four hours later, the supernatants were subjected to ELISA against SP and 5-HT (**Fig. 4a**). The results indicate that IL-33—but not SCF— stimulates mast cells to release SP and 5-HT.

**Figure 4.**
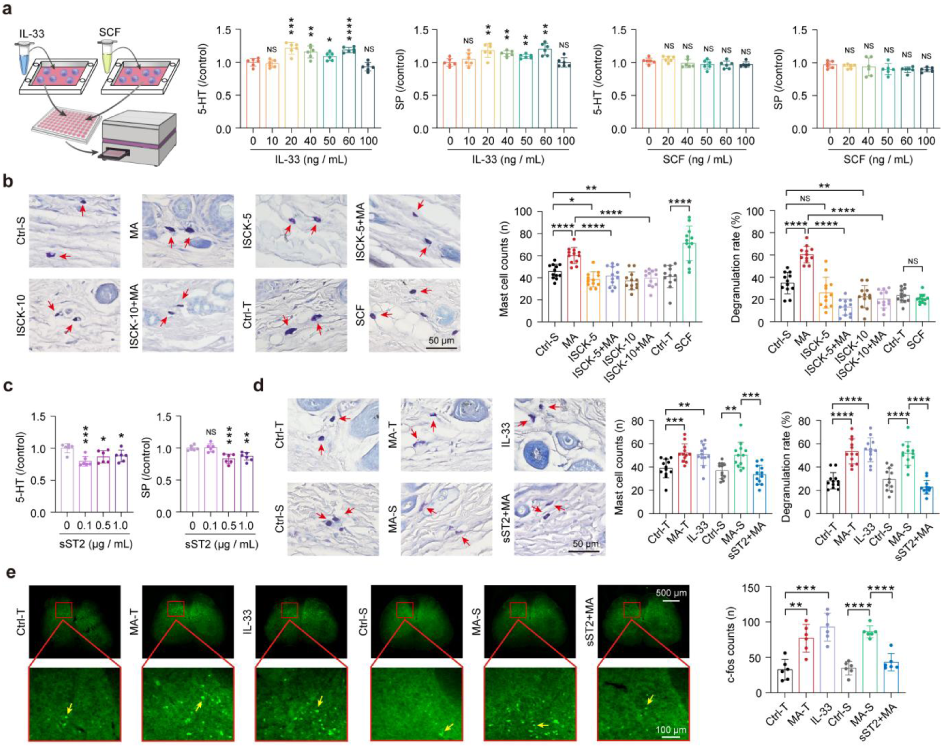
In the acupoint SCF and IL-33 regulate mast cell function and neural signal conduction. **a.** Schematic of IL-33 and SCF acting on mast cells. The level of SP and 5-HT in the cell supernatant was detected after the P815 cells were incubated with IL-33 and SCF solution for 4 hours (n = 6). **b.** Toluidine blue staining on the ST36 acupoint. Ctrl-S and Ctrl-T: PBS and 5% trehalose injection, respectively; ISCK-5 and ISCK-10: 5 µM and 10 µM ISCK03 injection, respectively; MA: manual acupuncture; SCF: SCF injection; ISCK-5+MA: 5 µM ISCK injection with MA; ISCK-10+MA: 10 µM ISCK injection with MA. Scale bars: 50 μm. The number of mast cells and the degranulation rate in the ST36 acupoint were statistically analyzed (n = 12). As each mouse has two ST36 acupoints, the number of the acupoint samples is twice the mice number. **c.** The level of SP and 5-HT in the cell supernatant under sST2 presence in the P815 cells (n = 6). d. same as b., IL-33: IL-33 injection; sST2+MA: sST2 injection with MA. **e.** Images (left) and quantification (right) of c-fos expression (green) in the dorsal horn of the lumbar spinal cords (n = 6). Scale bar: 100 μm.

SCF is well-recognized as a chemokine for mast cells in various conditions^39^. We tested whether SCF serves as the specific factor secreted by stretched fibroblasts that induces chemotaxis of P815 mast cells using the Transwell assays (**Extended Data Fig. 8**). The results suggested that the presence of the SCF facilitates mast cells in passing through the porous layer into the lower chamber; on the other hand, the presence of SCF inhibitor ISCK03 decreases the chemotaxis activity of mast cells, indicating that SCF is the only mast-cell chemokine present in the SSFC.

To further validate the *in vivo* role of SCF in acupuncture, we compared two conditions to assess whether SCF acts as the principal chemokine recruiting mast cells locally: (1) direct injection of SCF into the ST36 acupoint.to assess whether SCF can successfully recruit mast cells to the acupoints as the *in vitro* experiments suggested; (2) injection of the SCF inhibitor ISCK03 at the ST36 acupoint prior to acupuncture to evaluate the possibility that another mast cell attractant exists during acupuncture after the SCF is blocked.

Since SCF was dissolved in 5% trehalose and ISCK03 in PBS, these two conditions were evaluated separately. To examine the effect of SCF injection, two groups of mice were compared (n = 6): Ctrl-T, which received 5% trehalose injection; and SCF, which received SCF solution injection. To determine whether ISCK03 can block the effect of acupuncture, six groups of mice were compared: Ctrl-S (injection with PBS), MA, ISCK-5, ISCK-5+MA, ISCK-10, ISCK-10+MA. Here, ISCK-5 and ISCK-10 denote injections of the SCF inhibitor ISCK03 at 5µM and 10µM, respectively; group names containing “MA” indicate the application of acupuncture.

Accordingly, 5 µl of ISCK03 or SCF was injected into the ST36 acupoint of the respective groups, and acupuncture was applied to all “MA” groups 30 minutes after injection. After ninety minutes, we examined whether the quantity and degranulation rate of mast cells increases (**Fig. 4b**). Comparison of toluidine blue-stained tissue sections across groups revealed that SCF acts as the chemokine for mast cells in the ST36 acupoint, while ISCK03 effectively inhibited their chemotaxis during acupuncture. These results indicate that no other mast cell-attracting factor can substitute for SCF at the ST36 acupoint during acupuncture, establishing SCF as the principal mediator for mast cell recruitment.

As IL-33 is contained in SSFC and can induce mast cells to release SP and 5-HT, we evaluated whether IL-33 is the principal factor in the SSFC for mast cells releasing SP and 5-HT. SSFC with or without sST2 (0, 0.1, 0.5 or 1 μg/ml), the soluble receptor of IL-33, was separately added to the mast cell cultures and incubated for four hours. The corresponding supernatants were subjected to ELISA against SP and 5-HT (**Fig. 4c**). The analyses indicated that sST2 effectively blocks the SSFC’s capacity to induce mast cells releasing SP and 5-HT, suggesting that IL-33 is the principal molecule in the SSFC inducing mast cells to release SP and 5-HT during acupuncture.

We next investigated the contribution of IL-33 to acupuncture *in vivo*. First, we evaluated whether IL-33 induces mast cell degranulation in the acupoints as acupuncture does. Then, we injected sST2 in the acupoint to see whether it can prevent the acupuncture from taking effect. Since IL-33 is dissolved in 5% trehalose and sST2 is dissolved in PBS, we again prepared two independent sets of experiments for quantitative analysis. Thirty-six mice were intravenously injected with a virus that fluorescently labels expressed c-fos. Three weeks later, the mice were divided into six groups: Ctrl-T, MA-T, IL-33, Ctrl-S, MA-S, and sST2+MA. The first three were used to compare the effects of IL-33 and acupuncture on mast cells, whereas the latter three were used to verify that the observed effect is primarily mediated by IL-33. Here, MA-T and MA-S refer to injections of 5% trehalose or PBS, respectively, followed by acupuncture 30 minutes later. After ninety minutes, the mice were biopsied to collect the tissue sections of the acupoints and the lumbar dorsal horn, which provide insight on mast cell activities and the c-fos expression, respectively.

The tissue sections of the six groups were toluidine-blue-dyed to analyze the mast cell amounts and degranulation rates (**Fig. 4d**). The results indicated that both acupuncture and the injection of IL-33 into the acupoints increase the quantity and the degranulation rate of the mast cells. In addition, the sST2 blocks the effects of acupuncture, suggesting IL-33 as the main molecule in the acupoint causing mast cells degranulation.

Having established that fibroblasts respond to the acupuncture stimulation and convert these mechanical signals into the biochemical signals, SCF and IL-33, for mast cell activation, we further ensured that IL-33 triggers the neuronal signals propagating to spinal cords. The tissue sections from the dorsal horn of the lumbar spinal cords collected from the six groups were visualized by microscopy (**Fig. 4e**). The increasing expression of c-fos in the micrographs of the IL-33 group suggests that the presence of the IL-33 molecules in the acupoint can effectively excite the neuronal system.

## IL-33 releasing modulates endotoxemia

Acupuncture in the acupoint functions through SCF and IL-33, the molecules secreted from stretched fibroblasts during acupuncture manipulation. We therefore investigated how SCF and IL-33 contribute to disease control through the mouse disease model, LPS-induced endotoxemia.

To investigate the roles of IL-33 and SCF in acupuncture-mediated protection against endotoxemia, 104 mice were randomly assigned to four groups: Ctrl-T, MA-T, IL-33, and SCF. Mice in the Ctrl-T and MA-T groups received 5% trehalose injection, whereas those in the IL-33 and SCF groups received IL-33 and SCF injections, respectively. Mice in different groups were subjected to their corresponding treatment; then, all mice were subjected to LPS injection to induce endotoxemia. Serum inflammatory factors were measured in six mice per group after 90 minutes, and survival was monitored for the remaining animals over 168 hours (**Fig. 5a**). We find that the survival rates, from highest to lowest, are observed in the IL-33, MA-T, SCF, and Ctrl-T groups, respectively. In addition, the MA-T and IL-33 groups are significantly different from the Ctrl-T group with *p* < 0.05, respectively, while the SCF group is not significantly different from the Ctrl-T group.

**Figure 5.**
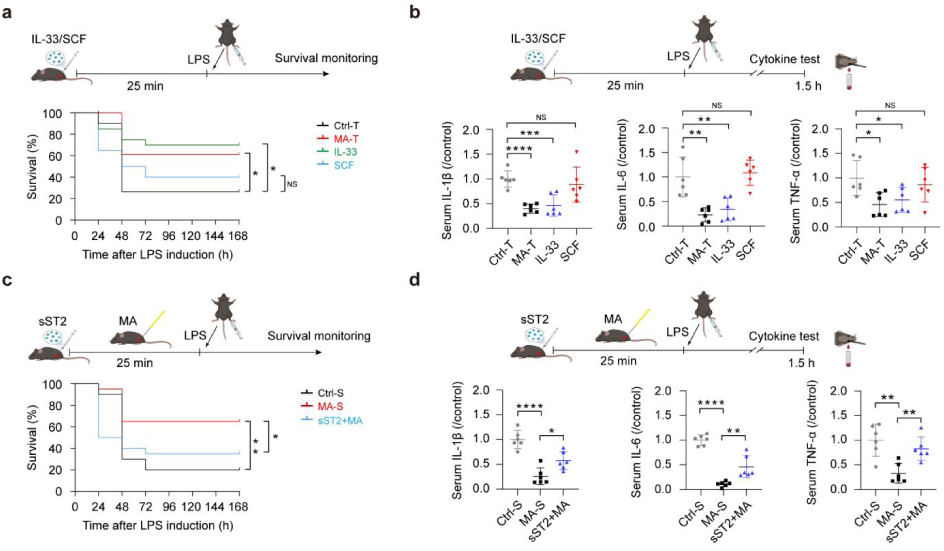
IL-33 mediates acupuncture effects against systemic inflammation. **a.** The survival rate of mice after specific treatment based on group for 25 minutes and intraperitoneal injection of LPS, recorded for 168 hours. Ctrl-T: 5% trehalose injection; MA-T: 5% trehalose injection with MA; IL-33: IL-33 injection; SCF: SCF injection (n = 20). **b.** Serum IL-1β/IL-6/TNF-α levels after intraperitoneal injection of LPS in mice (n = 6). **c.** Same as a. Ctrl-S: PBS injection; MA-S: PBS injection with MA; sST2+MA: sST2 injection with MA. **d.** Same as b. The significances in b and d were determined using Mantel-Cox log-rank test.

The serum samples of each group were separately subjected to ELISA for inflammatory factors (**Fig. 5b**). The results indicate that the MA-T and IL-33 groups exhibit low concentrations of IL-1β, IL-6 and TNF-α, while the SCF and Ctrl-T groups exhibit high concentration of these inflammatory factors. We find significant difference in IL-1β levels between the MA-T and Ctrl-T groups (*p* < 0.0001) and between the IL-33 and Ctrl-T groups (*p* < 0.001). Meanwhile, there is no significant difference between the SCF and Ctrl-T groups. There are also significant difference in IL-6 levels between the MA-T and Ctrl-T groups (*p* < 0.01) and between the IL-33 and Ctrl-T groups (*p* < 0.01). Furthermore, there is no significant difference between the SCF and Ctrl-T groups. On the other hand, significant differences exist between the MA-T and Ctrl-T groups and between the IL-33 and Ctrl-T groups (both with *p* < 0.05) in TNF-α, while there is no significant difference between the SCF and Ctrl-T groups.

We further tested whether the introduction of sST2 to the ST36 acupoint would interrupt the effectiveness of acupuncture through the same acupoint. Seventy-eight mice were randomly divided into three groups: Ctrl-S, MA-S, and sST2+MA. Mice in the sST2+MA group received sST2 injection at the ST36 acupoint, while mice in the Ctrl-S and MA-S groups were injected the same amount of PBS. Acupuncture was applied to the MA-S and sST2+MA groups; then, LPS was administered in all mice to induce inflammation. Consequently, survival rates of mice (n = 20) (**Fig. 5c**) and serum inflammatory factors (n = 6) (**Fig. 5d**) were compared across groups.

The survival rates again showed that acupuncture diminishes the LPS’ effect on mice inflammation and helps the survival of mice (*p* < 0.01) in the MA-S group; however, the presence of sST2 diminishes the effect of acupuncture on the MA-S group (*p* < 0.05). The concentrations of IL-1β, IL-6 and TNF-α in the serum also showed that the MA-S group has the lowest concentration of inflammatory factors (*p* < 0.0001, 0.0001 and 0.01, respectively); conversely, the sST2+MA group cannot maintain the low levels of inflammatory factors (*p* < 0.05, 0.01 and 0.01, respectively).

Taken together, we conclude that IL-33 is the primary molecule in the ST36 acupoint enabling effective acupuncture in the mouse model with LPS-induced endotoxemia.

Accordingly, we establish a working model of acupuncture as following (**Fig. 6**): During acupuncture operation, a needle is manually inserted into an acupoint and entwined the collagen fibers. Due to fiber entanglement, tension is transmitted through the network, stretching the fibroblasts anchored to it. In response to the tension, fibroblasts release SCF and IL-33 to the microenvironment. SCF has the chemotaxis capacity that attracts mast cells close by, while IL-33 induces mast cells to release neuronmodulatory molecules, such as SP and 5-HT. The neuromodulatory molecules further excite local neurons, evidenced by the c-fos expression in the dorsal horn of the lumbar spinal cord. Hence, local signals generated at the acupoint through acupuncture can be converted and propagated to exert physiological benefits. In the presence of the SCF inhibitor ISCK03 and the IL-33 inhibitor sST2, respectively, the ST36 acupuncture effects on preventing endotoxemia are eliminated. Hence, IL-33 and SCF are the principal mediators of anti-inflammatory effects triggered by fibroblasts stretching during acupuncture.

**Figure 6.**
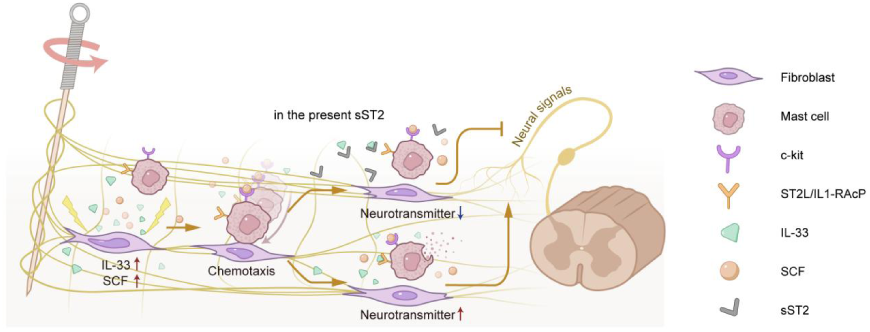
A working model of acupuncture. Schematic of the role of fibroblasts, IL-33 and SCF in acupuncture activation at the ST36 point.

## Discussion

In this study, we demonstrate that fibroblasts serve as the primary responders to the mechanical forces of acupuncture. We further identify stem cell factor (SCF) and interleukin-33 (IL-33) as pivotal molecules in the acupuncture response. Both factors are pre-stored within fibroblasts and are released immediately upon acupuncture stimulation, allowing for rapid responses to the mechanical environment. SCF plays a crucial role in recruiting mast cells, whereas IL-33 directs mast cells to release the neuromodulatory factors. Consequently, acupuncture mediates selective nerve activation through paracrine signaling from fibroblasts, facilitating the transmission of acupuncture-related information through mast cells to the nervous system.

Traditionally recognized as a regional alarm molecule, IL-33 is released in response to stimuli, recruiting macrophages and driving local immune responses. However, our findings reveal that IL-33 is central to the therapeutic effects of acupuncture, promoting mast cells to release neuromodulatory factors in the acupuncture area. Notably, we observed that the injection of an appropriate dose of IL-33 into the ST36 acupoint—without the need for needle manipulation—can replicate the protective effects against endotoxemia in mice.

In the 1930s, Swiss physician Paul Niehans’s use of sheep embryo cells in human treatment was rooted in anecdotal observation rather than mechanistic understanding. Those early efforts paved the way for today’s rigorously defined cell-based therapies, such as CAR-T immunotherapy. Similarly, acupuncture has been practiced for centuries, and its therapeutic efficacy has long been recognized. Our findings serve as a bridge between the enigmatic “black box” and the physiological effects accompanying well-defined cellular regulatory mechanisms. As its scientific basis continues to emerge from this “black box,” acupuncture is poised to evolve rapidly into a pillar of modern medicine—advancing both human health and our understanding of the body. By bridging traditional medical practices with contemporary scientific frameworks, our work demonstrates that traditional medicine—rooted in generations of empirical knowledge and cultural heritage—can be rigorously examined and meaningfully integrated into modern biomedical science.

## Methods (including separate data and code availability statements)

### Mice maintenance

All C57/BL6J male mice used in this study were purchased from Jinan Pengyue Laboratory Animal Breeding Co., Ltd, Shandong, China. The mice were acclimatized in a controlled environment (12-hours light/dark cycle, 22 ± 2°C, 55 ± 5% relative humidity) with free access to food and water for one week prior to the experiments. All the experimental procedures strictly adhered to the “Guide for the Care and Use of Laboratory Animals” and were approved by the Experimental Animal Ethics Committee of Shandong University of Traditional Chinese Medicine (Approval No. SDUTCM20241128001).

### LPS induction of endotoxemia in mice

Male mice were used since they have greater Lipopolysaccharide (LPS) sensitivity compared to females^12^. LPS from Escherichia coli strain 0111:B4 (L2630, Sigma-Aldrich) was formulated as a 10 mg/ml solution in sterile, pyrogen-free PBS (Gibco Life Technologies) and stored at 4°C. Mice were intraperitoneally injected with a LD_80_ (80% lethality rate) dose of LPS (8 mg/kg for C57BL/6J male mice). Mouse survival rate was monitored as previously described^13^. Whole blood was collected from mouse eyeballs 1.5 hours after LPS injection.

### Manual acupuncture

Mice were anaesthetized with isoflurane (0.5 - 1.5%) and the body temperature was maintained using a heating pad. Manual acupuncture (MA) was conducted with stainless-steel acupuncture needles (0.25 × 13 mm). Acupuncture was administered at hindlimb ST36 (Zusanli) acupoints, located about 4 mm down from the knee joint and about 2 mm lateral to the anterior tubercle of the tibia. Acupuncture needles kept inserting in the acupoints, received bidirectional manipulation (180° rotation amplitude at 2 Hz) once every 5 minutes for 1-minute duration, and remained still for 4 minutes for a total of 5 times. The total treatment duration was 30 minutes.

### Subcutaneous injection in acupoint area and tail vein injection

For fibroblast depletion, C57/BL6J mice (5 weeks) were anaesthetized with an isofluorane/air mix (2% for initial induction and 1.5% for maintenance). A mixture of two adeno-associated viruses (AAVs) was administered via subcutaneous injection at the ST36 acupoints using a Hamilton syringe with 30G needle, connected to a microinjection pump. The viral cocktail consisted of AAV5-Fsp1-Cre-WPRE (≥ 2×10^12^ vg/mL, for fibroblast-specific labeling) and rAAV5-CMV-DIO-taCasp3-TEVp (≥ 2×10^12^ vg/mL, for inducible apoptosis), combined at a 1:2 ratio in PBS. Injection volumes of 300, 600, 1200, and 1500 nL were tested to establish optimal depletion efficiency.

For collagenase, IL-33, sST2, SCF, or ISCK03 injection, C57/BL6J mice (8 weeks) were anaesthetized as previously described. Each ST36 acupoint (both sides) received 5 μL of type I collagenase (5 mg/mL; 17100-017, Gibco), IL-33 (4 ng/μL; 210-33, Thermo Fisher), soluble ST2 (sST2; 30 ng/μL; 1004-MR, R&D), SCF (2 ng/μL; 250-03, Thermo Fisher), or ISCK03 (5 μM; HY-101443, MCE).

For newly expressed c-fos labeling, C57BL/6J mice (5 weeks) received intravenous injections of rAAV2-cFos-EYFP (2×10¹¹ viral genomes/mouse in 100 μL total volume) via the lateral tail vein.

### Cytokine measurement

Collected cell-culture media was sequentially centrifuged (300×g for 5 minutes and 10,000×g for 20 minutes) at 4°C to remove debris. The level of 5-HT, SP, NGF, M-CSF, CXCL1, IL-33, and SCF were analyzed using ELISA kits (E-EL-0033c, E-EL-0067c, E-EL-M0815, E-MSEL-M0105, E-EL-M0018, E-EL-M2642c, and E-EL-M0636c, respectively) from Elabscience, following manufacturer’s instructions.

For *in vivo* serum cytokine analyses, blood was collected from mouse eyeballs after 1.5 hours of LPS injection, allowed to clot for 2 hours at room temperature, and then centrifuged at 4000 rpm for 10 minutes at 4°C. The serum was collected and stored at -80°C before use. Levels of IL-1β and TNF-α were analyses by ABplex Mouse 2-Plex Custom Panel (RK04380, ABclonal). Level of IL-6 were analyzed by ELISA kit (E-EL-M0044, Elabscience), following manufacturer’s instructions.

### Immunohistochemistry

Three days before the experiment, the hair around the mice’s ST36 acupoints was shaved. Then, the mice were transcardially perfused with PBS and 4% paraformalaldehyde (PFA; pH 7.4) sequentially. The tissue centered by the ST36 acupoint area (5 mm × 5 mm × 2 mm) and the spinal cord at lumbar segments were dissected and post-fixed in 4% PFA overnight at 4°C. Spinal cord were cryopreserved in 30% sucrose in PBS overnight and then embedded in Tissue-Tek OCT compound (NEG-50, Epredia). For immunofluorescence staining of spinal cords, 30 μm thick sections were made with a cryostat and mounted on SuperFrost Plus slides (CM1950, LEICA). The sections of ST36 area were blocked with 0.3% Triton X-100 (Sigma-Aldrich) plus 5% goat serum (SL038, Soarbio) in PBS for 1 hour and then incubated with rabbit anti-fibronectin (1:300, RT1224, HUABIO) overnight at 4°C. After washing with 0.3% of Triton X-100 in PBS, the sections were incubated with secondary antibody goat anti-rabbit Alexa Fluor 488 (1:1000; ab150077, Abcam) for 1.5 hours at room temperature, and DAPI nuclear counterstain (5 μg/mL, C1002, Beyotime) for 10 minutes at room temperature. Images were acquired using a BZ-X800 fluorescence microscope (KEYENCE).

Tissue specimens of ST36 acupoints were processed through an ethanol gradient for dehydration, embedded in paraffin, and sectioned at 5 μm thickness using a microtome (HistoCoreAUTOCUT, LEICA). To delineate collagen fiber distribution in ST36 acupoint, tissue sections were stained using Masson’s trichrome technique according to the manufacturer’s protocol. To quantify mast cell populations and assess morphological features, cationic toluidine blue staining was performed according to manufacturer protocols. Mast cells were examined under 400× magnification light microscopy (10 × 40). For each section, twenty randomly selected fields were analyzed to determine total mast cell counts and the number of degranulated mast cells. Degranulation rates were calculated as: Degranulation Rate (%) = (Number of Degranulated Mast Cells / Total Mast Cells) × 100%.

### Multiplex immunohistochemistry and multichannel imaging

Deparaffinization of tissue sections of ST36 acupoints was done through xylenes and rehydration through decreasing graded alcohol. AR6 buffer (Akoya Biosciences) was used for antigen retrieval in a microwave oven. Endogenous peroxidase was inactivated by incubation in 3% H_2_O_2_ for 10 min. Multiplex immunohistochemistry was performed by several rounds of staining, each including a protein block with 1% BSA followed by primary antibody and corresponding secondary horseradish peroxidase-conjugated antibody against rabbit immunoglobulins (Akoya). The slides were then incubated in different Opal fluorophore (NEL861001KT, Akoya), 1:100 diluted in 1X Plus Amplification Diluent (Akoya). After tyramide signal amplification and covalent linkage of the individual Opal fluorophores to the relevant epitope, the primary and secondary antibodies were removed via antigen retrieval as previously mentioned and the next cycle of immunostaining was initiated. The sequence of primary antibody/Opal fluorophore was rabbit anti-c-kit (1:500, GB113799, Servicebio)/Opal 620, rabbit anti-vimentin (1:1000, ab92547, Abcam)/Opal 570 and rabbit anti-collagen I (1:500, ab270993, Abcam)/Opal 690. All slides were counterstained with spectral DAPI (Akoya) and mounted with anti-fade fluorescence mounting medium (ab104135, Abcam). Multichannel imaging was performed on a PANNORAMIC SCAN II Imaging System (3Dhistech, Hungary). Slides were imaged at ×200 magnification.

### Cell culture

The murine mastocytoma P815 cell line (obtained from the Chinese Academy of Sciences Cell Bank) was maintained in RPMI 1640 medium (Gibco) supplemented with 10% fetal bovine serum (FBS; Corning) at 37°C under 5% CO₂. Primary skin fibroblasts were isolated from neonatal mice: After 5-minute disinfection in 75% ethanol and washes twice with PBS containing 1% penicillin-streptomycin, skin was incised using ophthalmic scissors at the neck, limbs and abdomen (avoiding peritoneal puncture). Full-thickness skin was dissected cervico-caudally, washed twice in antibiotic-supplemented PBS, and subcutaneous fat was gently removed. Minced tissue underwent mechanical/enzymatic dissociation for 25 minutes in a tissue homogenizer tube, followed by digestion termination. Then, the cell suspension was sequentially filtered through a 70 μm mesh. After repeated centrifugation (300×g, 5 minutes, 4°C) and re-suspension (with 5% BSA solution containing 8% penicillin-streptomycin), cells were cultured in DMEM (Corning) with 10% FBS, 1% L-glutamine (Gibco) and 1% penicillin-streptomycin (Gibco) at 37°C in 10% CO₂.

### Identification of skin primary fibroblasts

The primary fibroblasts were identified by fluorescence microscopy. Cell samples were seeded into a 48-well plate at an initial density of 3×10^4^ cells/well. After 2 days of cultivation, cells were fixed by prewarmed 4% paraformaldehyde (Sigma-Aldrich) in PBS for 15 minutes, permeabilized by 0.25% Triton X-100 in PBS for 10 minutes, and blocked by 5% BSA in PBS for 1 hour at room temperature. Afterwards, cells were incubated with rabbit anti-vimentin (ab92547, Abcam) overnight at 4°C and washed 3 times by PBS. Then, Alexa 568-labeled goat anti-rabbit IgG (ab175471, Abcam) was applied as secondary antibody for 1 hour at room temperature before 3 times PBS washes. For DNA staining, cells were incubated with 0.1 μg/ml Hoechst 33342 (BMD00062, Sigma) at the same time as the secondary antibody. The cell images were acquired by a BZ-X800 fluorescence microscope (Keyence).

### Lentivirus transfection

HEK293TA cells (Hanbio, Wuhan, China) were cultured in DMEM, supplemented with 10% FBS 1 day before lentivirus preparation. Lenti-GFP plasmids (Hanbio) were transfected into HEK293TA cells using LentiFit transfection reagent (Hanbio) according to the manufacturer’s protocol. Media containing lentiviral particles were harvested 48 hours post-transfection, aliquoted, and stored at -80°C. For GFP labeling, fibroblasts were exposed to lentiviral media for 3 days to achieve stable GFP expression.

### Mechanical stimulation of the cells

Cells were stimulated with continuous cyclic stretching (stretch ratio: 5% or 10%, stretch frequency: 0-2 Hz,) using a cell-stretching system (ShellPa Pro, Menicon Life Science), fitted with polydimethylsiloxane (PDMS) stretch chambers. Prior to cell seeding, stretch chambers were sequentially coated with poly-L-lysine (PLL; P4832, Sigma), followed by collagen solution (100 μg/mL; 07005, Stemcell). P815 cells were seeded at 2.0×10⁵ cells/mL. For primary fibroblasts, chambers received extra fibronectin coating (20 μg/mL; F0895, Sigma) before seeding at 5.0×10⁴ cells/mL.

### Adhesion of mast cells

Cell adhesion experiments were performed on 35-mm dishes with 4 types of surface treatments: untreated control, poly-L-lysine (PLL), collagen (100 μg/mL), or fibronectin (20 μg/mL). Mast cells (2.0×10⁵ cells/mL) were pre-stained with Hoechst 33342, seeded onto coated dishes, and incubated for 4 hours at 37°C with 5% CO₂. Using live-cell imaging under controlled conditions (37°C and 5% CO₂), cells underwent microfluidic perfusion at 1.489 mL/minute for 4-times complete media exchanges. Adherent cells were quantified from 15 randomized tile-scan fields (5×5 images/field), with adhesion efficiency calculated as: Adhesion rate (%) = Dish surface area × Number of adherent cells / Total cells seeded / Imaged area × 100%.

### Immunofluorescence (IF) staining of cell

Cells were washed thrice with pre-warmed HBSS, fixed in 4% paraformaldehyde (15 minutes, room temperature), and permeabilized with 0.1% Triton X-100 (10 minutes, room temperature). Following PBS washes, samples were blocked with 1% of BSA plus 0.1% Tween-20 in PBS for 1 hour at room temperature, and then incubated with primary antibodies overnight at 4°C: rabbit anti-vimentin (1:1000; ab92547, Abcam), rabbit anti-fibronectin (1:300; RT1224, HUABIO), rabbit anti-IL-33 (1:500; ab187060, Abcam), or rabbit anti-SCF (1:250; 26582-1-AP, Proteintech). After washing with PBS, samples were incubated with secondary antibody goat anti-rabbit Alexa Fluor 488 (1:1000; ab150077, Abcam) or goat anti-rabbit Alexa Fluor 568 (1:1000; ab175471, Abcam) for 1 hour at room temperature, followed by phalloidin-647 (1:1000; ab176759, Abcam) for 30 minutes and Hoechst 33342 nuclear counterstain for 10 minutes at room temperature. Final PBS washes preceded fluorescence microscopy imaging.

### Modulation of P815 cells by mechanically stimulated fibroblasts

Primary skin fibroblasts (5.0×10⁴ cells/mL) were seeded onto coated stretch chambers and cultured for 48 hours at 37°C with 10% CO₂. After media replacement, samples were divided into control and stretched groups, with the latter subjected to 5% cyclic strain at 0 Hz (static), 0.5 Hz, or 2 Hz for 30 minutes. Conditioned media was sequentially collected through centrifugation (300×g, 5 minutes, 4°C and 10,000×g, 20 minutes, 4°C) to remove debris. P815 cells (2.0×10⁵ cells/mL) were then cultured in this media for 4 hours. In parallel co-cultures, P815 cells (2.0×10⁵ cells/mL) were directly seeded onto fibroblasts and stretched under identical parameters. Post-stimulation supernatants were collected after centrifugation as previously mentioned for 5-HT and SP quantification.

### Mast cell regulations by IL-33, SCF and sST2

P815 cells were stimulated for 4 hours with recombinant IL-33 (10-100 ng/mL), SCF (20-100 ng/mL), or soluble ST2 (sST2; 100-1000 ng/mL). Supernatants were cleared by centrifugation (10,000×g, 20 minutes) for 5-HT and SP measurements.

### Transwell chemotaxis experiments

Transwell inserts (5 μm pores) were coated with fibronectin on the underside. Inserts were placed in 24-well plates with one of the chemoattractant solutions (650 μL) in the lower chamber. The chemoattractant solutions include complete fibroblast medium, unstretched fibroblast-conditioned medium, conditioned medium from fibroblasts subjected to 5% static strain (0 Hz) for 30 minutes, and SCF solution. The P815 cells (8×10⁵ cells/mL, 100 μL) with or without the presence of ISCK03 were loaded to upper chambers. After assembled, the Transwell device were incubated at 37°C with 10% CO₂ for 8, 12, or 16 hours (for examining the effects; and later on, merely incubated for 12 hours). Non-migrated cells were removed by cotton swab abrasion. Migrated cells on the membrane underside were fixed (4% PFA, 10 minutes), stained with 0.1% crystal violet (20 minutes), PBS-washed, and quantified by counting 10 random 20× fields per insert using Image J 1.53a.

### Statistics

The statistics followed by Liu, S. *et al*.^12^ Briefly, results are expressesd as mean ± sd. Statistical analyses were done using GraphPad Prism v9.5. Image analyses were done using Image J 1.53a. Survival rates are expressed using Kaplan-Meier curves, and the comparison among survival curves were performed with a Mantel-Cox log-rank test. Other data were analysed with two-side Student’s unpaired *t*-test.

## Data availability statement

The data that support the findings of this study are available from the corresponding author upon request.

## Acknowledgements

This work was financially supported by National Key R&D Program of China (NO. 2022YFC3500405 to YDT), China Postdoctoral Science Foundation (NO. 2024MM761884 to XQX), Shandong Province Traditional Chinese Medicine Science and Technology Project (NO. M20241713 to XQX), Shandong Provincial Natural Science Foundation Innovation and Development Joint Funding (NO. ZR2022LZY023 to TL), and Taishan Scholar Foundation of Shandong Province (NO. tstp20221125 to YDT). The authors are thankful for Experimental Center, Shandong University of Traditional Chinese Medicine.

## Contributions

YDT and XQX designed the experiments, analyzed the data, wrote the original paper draft. TL reviewed the paper draft. MY conducted bioinformatic and partial imaging data analysis. XQX, RW, WS, JP, DL conducted the experiments. All data were generated inhouse, and no paper mill was used. All authors agree to be accountable for all aspects of work ensuring integrity and accuracy.

## Corresponding author

Correspondence to Yiider Tseng.

## Ethics declarations

The authors declare that they have no conflict of interest. The authors declare that there is no known competing financial interests or personal relationships that could have appeared to influence the work in this report.

## Extended data figures

### Interaction between acupuncture needle and the tissue of ST36 acupoint

We examined what happens when the acupuncture needle was inserted into the ST36 acupoint using B-scan ultrasound (**Extended Data Fig. 1a**). The ultrasonographs shows needle wrapped by the white fibers inside the acupoint tissue. After the needle was withdrawn from the acupoint, several white fibers remained attached. The Masson’s trichronme stain was applied to the tissue sections of the ST36 acupoint (**Extended Data Fig. 1b**) and the graphs display abundant blue fibers. Since collagen fibers are known to exhibit blue color with this stain, we considered the fibers to be collagen. The graphs show that the dermis region exhibits markedly denser and more abundant collagen fibers than the subcutaneous connective tissue. To confirm the fibers contain collagen, we further conducted the Western blot against the material attached to the acupuncture needle after the needle was inserted and rotated in the acupoint (**Extended Data Fig. 1c**). The material was homogenized before conducting the Western blotting against collagen I antibody, indicating the presence of collagen I in the needle-adherent substrate.

**Extended Data Fig. 1.**
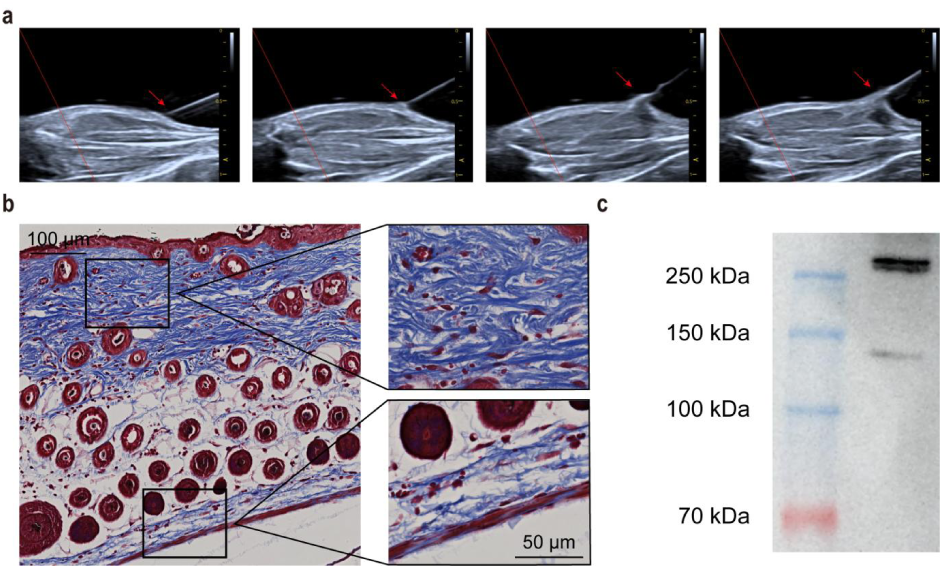
Needle manipulation in the ST36 acupoint. **a**. B-scan ultrasonographs display that relation between the inserted needle and the acupoint’s tissue (indicated by red arrows). **b.** Masson’s trichrome staining of paraffin-embedded ST36 (Zusanli) acupoint sections in mice. Scale bar: 100 µm (left). The square images are magnified (right). Scale bar: 50 µm. **c.** Material adhering to the needle was analyzed by Western blot using antibody against collagen I.

### Acupoint-fibroblast depletion and labelling through two adeno-associated viruses

To determine whether dermal fibroblasts inside the ST36 acupoint are essential for acupuncture effecacy, the ST36 acupoint were injected with PBS containing two adeno-associated viruses (AAVs), mixed at a 1:2 ratio. The former one was AAV5-fsp1-cre-WPRE-PA, which fluorescently labels fibroblasts, and the latter one was rAAV5-CMV-DIO-taCasp3-TEVpWPRE-hGH polyA, which induces fibroblast apoptosis. The AAVs solution was injected at ST36 at three volumes, 600, 1200, and 1500nL, while the control group received 1500 nL of PBS. Three weeks later, biopsies were conducted on the tissue of the ST36 acupoint (**Extended Data Fig. 2**). The tissue sections revealed the acupoint injected with more than 1200 nL of AAV solution have markedly reduced fibroblast fluorescence, compared with controls, confirming effective local depletion after injecting 1200 nL AAVs.

**Extended Data Fig. 2.**
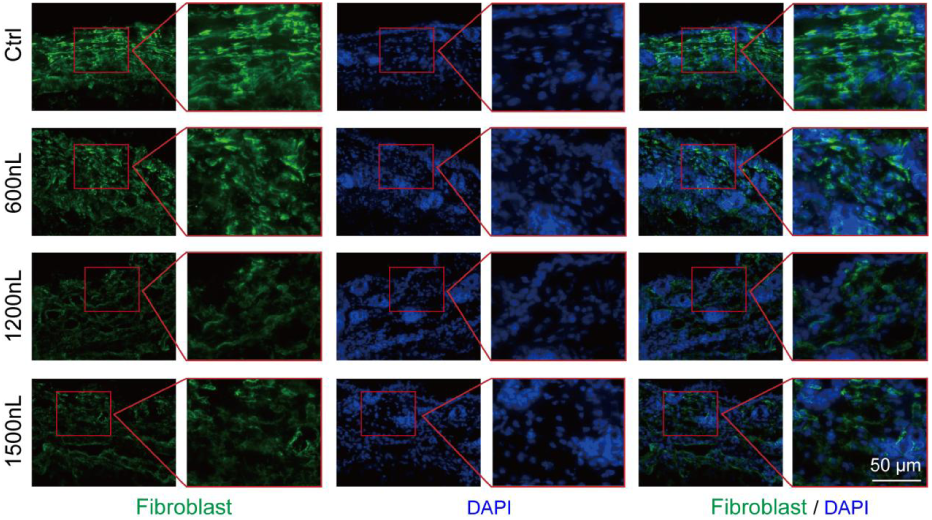
Fibroblast depletion through the AAV injections at the ST36 acupoint. AAV was injected at ST36 in three volumes (600, 1200, and 1500 nL), while control group received 1500 nL of PBS. Biopsies of the ST36 acupoint were conducted three weeks after injection. Fibroblasts were labeled with green color. Scale bar: 50 µm.

### Documentation of dorsal horn c-fos expression through an adeno-virus labelling

To accurately trace the transmission of mechanical force signals generated by acupuncture from the acupoint to spinal cord in real time, mice received tail vein intravenous injection of rAAV2-cFos-EYFP (≥ 5×10^12^ vg/mL) at a dose of 2×10^11^ vg/mL per mouse (**Extended Data Fig. 3a**). Three weeks after injection, spinal cords were harvested at the lumbar level 90 minutes post-stimulation (manual acupuncture at the ST36 acupoint) and cryo-sectioned for fluorescence microscopy (**Extended Data Fig. 3b**). Robust c-fos expression (green) was observed in the dorsal horn, compared to the control, indicating neural activation following acupuncture.

**Extended Data Fig. 3.**
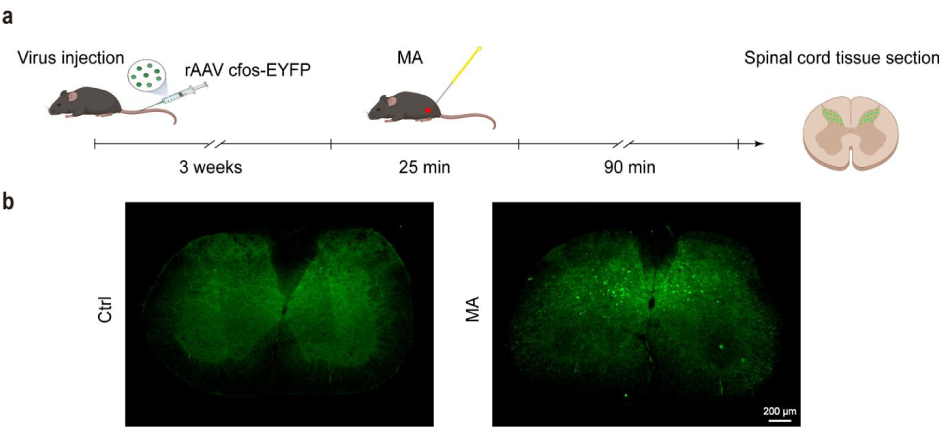
Fluorescently labelled c-fos in the dosal horn of the lumbar spinal cords. **a.** The procedure for the introduction of c-fos labeling to the dosal horn of the lumbar spinal cords. **b.** The fluorescent micrographs of the dorsal horn indicate the expression level of c-fos: the dorsal horn of mouse with (right) and without (left) prior acupuncture stimulation.

### Acupuncture stimuli cause the re-distributions of cells in the ST36 acupoint

To access the dynamic distributions in fibroblasts and mast cells within the ST36 acupoint after acupuncture stimulation, the acupoint tissues were collected, paraffin-embedded, and sectionized 90 minutes after acupuncture for multiplex immuno-fluorescence staining (**Extended Data Fig. 4**). The interactions and morphological changes of fibroblasts and mast cells were examined through antibodies against collagen I (red), fibroblast marker vimentin (green) and mast cell marker c-kit (yellow). Comparing with tissue sections without acupuncture, tissues subjected to acupuncture stimuli exhibited increased mast cell density, cellular hypertrophy, and dispersed fluorescence signaling. Furthermore, acupuncture significantly reduced the intercellular distance between fibroblasts and mast cells.

**Extended Data Fig. 4.**
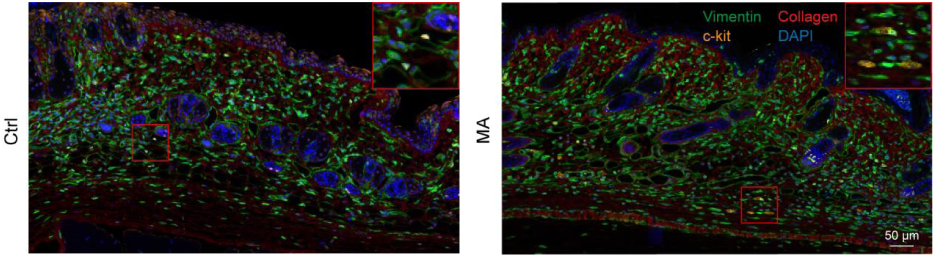
Markers of fluorescence immunostaining against fibroblast, mast cell and collagen I on the ST36 tissue section with and without acupuncture stimuli. The tissue section without (left) and with (right) acupuncture stimuli. Through antibody labelling, collagen I, fibroblast marker vimentin and mast cell marker c-kit colored in red, green and yellow, respectively. Scale bar: 50 µm.

### The attachments of fibronectin to the fibroblasts *in vitro* and *in vivo*

To determine whether dermal fibroblasts are coated with fibronectin that facilitates fibroblasts-mast cell contact, immunofluorescence staining was performed on both primary murine dermal fibroblasts *in vitro* (cultivated for 2 days) (**Extended Data Fig. 5a**) and in the tissue section of the ST36 acupoint (**Extended Data Fig. 5b**). The micrographs revealed that abundant fibronectin appear on the surface of primary murine dermal fibroblasts (red) and throughout fibroblasts in the ST36 acupoint tissue sections (green).

**Extended Data Fig. 5.**
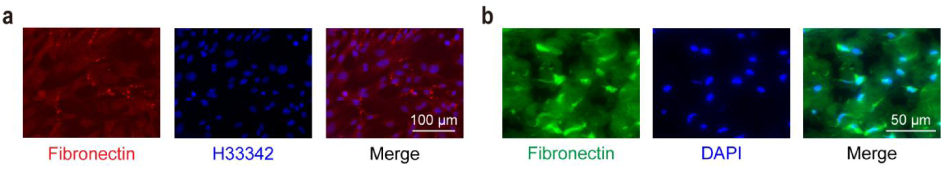
Fluorescence immunostaining against fibronectin *in vivo* and *in vitro*. **a.** From left to right, the micrographs of 2-day primary murine dermal fibroblast culture are labeled with fibronectin (red), nuclei (blue) and merged colors. Scale bar: 100 µm. **b.** From left to right, the micrographs of ST36 tissue section are labeled by fibronectin (green), nuclei (blue) and merged colors. Scale bar: 50 µm.

### The comparison of fibroblast morphology change between *in vivo* and *in vitro*

It has been shown that acupuncture stimuli increase fibroblast size^38^. To establish optimal baseline for the mechanical stimulation parameters in murine primary dermal fibroblasts, we exogenously introduced GFP into fibroblasts via lentiviral transduction. Then, the GFP-expressing fibroblasts were cultured on the fibronectin-coated stretching device and subjected to a one-hour mechanical stimulation at 5% or 10% strain amplitudes using a cell-stretching system (**Extended Data Fig. 6**). The 5% stretch induced noticeable cellular hypertrophy to fibroblasts, whereas the 10% stretch caused cell partially detachment from the substrate. Hence, the morphological changes in the 5% stretched fibroblasts more closely resembled those of fibroblasts subjected to the acupuncture in the tissue.

**Extended Data Fig. 6.**
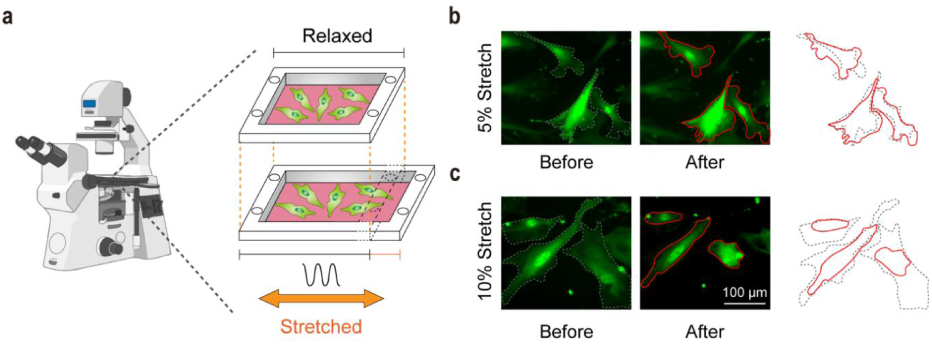
The GFP-labelled fibroblasts before and after being subjected to stretching. **a.** A polydimethylsiloxane (PDMS) cell culture device is utilized to apply stretching treatment. **b.** The fibroblasts before (left) and after (middle) subjected to 5% stretching. Cell morphology before (dash line) and after (solid line) stretching are overlaid for comparison (right). **c.** Fibroblasts before (left) and after (middle) subjected to 10% stretching. Cell morphology before (dash line) and after (solid line) stretching are overlaid for comparison (right). Scale bar: 100 µm.

### Bioinformatic studies of fibroblasts under stretching

We probed the transcriptional responses of primary murine dermal fibroblasts with mechanical stimulation. Primary murine dermal fibroblasts were cultured on fibronectin-coated stretch chambers for 48 hours and subjected to cyclic mechanical stimulation (5% amplitude and 1 Hz frequency for 30 minutes) using a cell-stretching system, with unstimulated cells serving as controls. Following mechanical stimulation, cells were immediately collected for RNA extraction and subsequent transcriptome sequencing (**Extended Data Fig. 7**). As shown in Extended Data Fig. 7a, no genes met the significance threshold under the stringent criterion (adjusted *p* < 0.05). Nevertheless, a permissive cutoff (*p* < 0.05) revealed 1567 differentially expressed genes, most of which were up-regulated following stretching. Gene Ontology (GO) and Kyoto Encyclopedia of Genes and Genomes (KEGG) pathway enrichment analyses through the STRING database indicated that these genes were predominantly associated with protein translation and ribosome biogenesis processes. Consistently, single-sample gene set enrichment analysis (ssGSEA) demonstrated significantly elevated enrichment scores for both ribosome and translation gene sets in stretched fibroblasts, suggesting that fibroblasts under short-term mechanical stretching remain engaged in the stages of ribosome biogenesis and protein translation rather than progressing to produce new secretory proteins.

**Extended Data Fig. 7.**
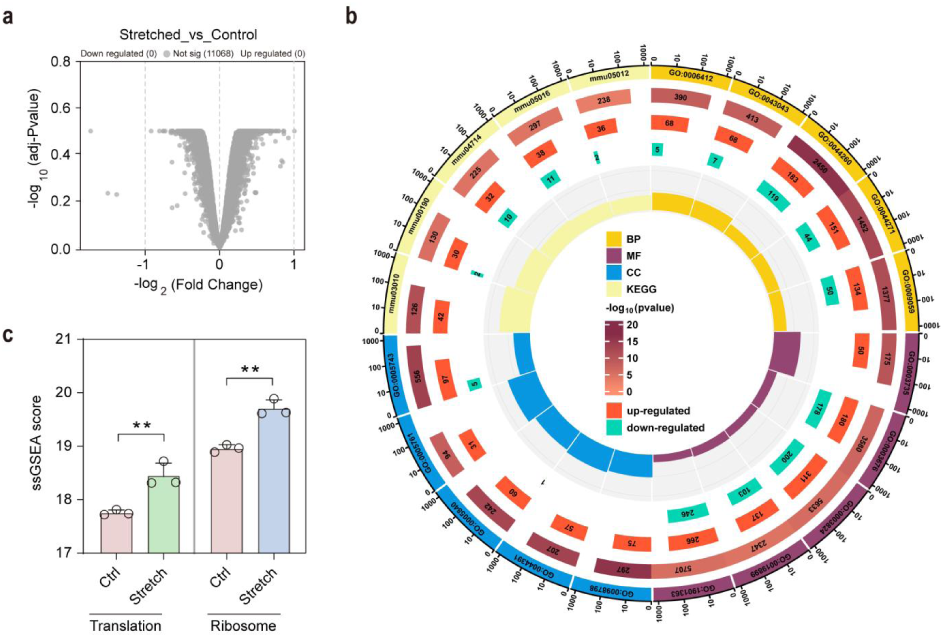
Bioinformatic study of stretched fibroblasts. **a.** Volcano plot showing no differentially expressed genes under the stringent threshold (adjusted *p* < 0.05). **b.** Circular Gene Ontology (GO) and Kyoto Encyclopedia of Genes and Genomes (KEGG) plot depicting the enriched terms with identified genes under permissive cutoff, alongside the numbers of respective up- and down-regulated genes. **c.** Bar plot illustrating single-sample gene set enrichment analysis (ssGSEA) enrichment scores for the ribosome and translation gene sets in stretched fibroblasts compared with controls. ** denote *p* < 0.01.

### The chemoattractant capacity of SCF from stretched fibroblasts

To determine the chemoattractant potential of SCF towards mast cells, we added varying concentrations of SCF solution (0, 10, 20, and 50 ng/mL) to the lower chambers of the Transwell plates and seeded P815 mast cells (8×10⁴ cells) in the upper chambers. After 12 hours of incubation, the upper membrane surfaces were stained with crystal violet and transmigrated mast cells were quantified (**Extended Data Fig. 8a**). SCF at 10 and 20 ng/mL significantly increased mast cell migration across the membranes, indicating that SCF promotes mast cell chemotaxis.

To confirm the specificity of SCF-mediated chemotaxis, we supplemented the supernatant from the stretched fibroblast culture (SSFC) in the lower chambers of Transwell, and seeded P815 mast cells (8×10⁴ cells) with 0, 1, 5, 10, or 20 μM of ISCK03, a c-kit receptor inhibitor, in the upper chambers to reassess mast cell migration using the Transwell assays (**Extended Data Fig. 8b**). After 12 hours of incubation, the upper membrane surfaces were stained with crystal violet and transmigrated mast cells were quantified. ISCK03 treatment (1, 5, and 10 μM) significantly reduced mast cell migration towards SSFC compared to the control group (0 μM ISCK03), demonstrating effective inhibition of SCF-mediated mast cell chemotaxis.

**Extended Data Fig. 8.**
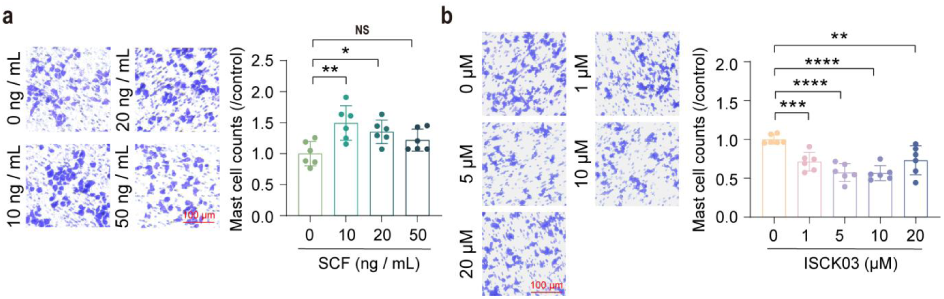
The evaluation of chemoattractant potential of mast cells in the presence of SCF. **a.** The analysis of the Transwell experiments of mast cells in the presence of SCF as the chemokine. Images of the cells after chemotaxis (left). The statistical analysis (right) (n = 6). **b.** The analysis of the Transwell experiments of mast cells in the presence of the conditioned medium from mechanically-stimulated fibroblasts with ISCK03. Images of the cells after chemotaxis (left). Scale bars: 100 µm. The statistical analysis (right) (n = 6). *, **, ***, **** denote *p* < 0.05, 0.01, 0.001, 0.0001, respectively. NS denotes non-significant, *p* > 0.05.

