## Supplementary Table 1 for "Decoding the fibroblast/mast cell signaling pathway of acupuncture"

### Supplementary Table 1: Statistical analyses

| Figure | Variables analysed (Two-side Student's unpaired *t*-test) | P-value | t | df |
|---|---|---|---|---|
| Fig. 1c (IL-1β) | MA *vs.* Ctrl | 0.0043 | 3.668 | 10 |
| Fig. 1c (IL-1β) | $FB^{dep}$+MA *vs.* $FB^{dep}$ | 0.0534 | 2.190 | 10 |
| Fig. 1c (IL-1β) | $FB^{dep}$+MA *vs.* MA | 0.0064 | 3.431 | 10 |
| Fig. 1c (IL-6) | MA *vs.* Ctrl | 0.0009 | 4.650 | 10 |
| Fig. 1c (IL-6) | $FB^{dep}$+MA *vs.* $FB^{dep}$ | 0.0835 | 1.922 | 10 |
| Fig. 1c (IL-6) | $FB^{dep}$+MA *vs.* MA | 0.0180 | 2.826 | 10 |
| Fig. 1c (TNF-α) | MA *vs.* Ctrl | 0.0038 | 3.749 | 10 |
| Fig. 1c (TNF-α) | $FB^{dep}$+MA *vs.* $FB^{dep}$ | 0.4138 | 0.8526 | 10 |
| Fig. 1c (TNF-α) | $FB^{dep}$+MA *vs.* MA | 0.0013 | 4.394 | 10 |
| Fig. 1e | MA *vs.* Ctrl | ＜0.0001 | 9.353 | 10 |
| Fig. 1e | $FB^{dep}$+MA *vs.* $FB^{dep}$ | 0.8446 | 0.2012 | 10 |
| Fig. 1e | $FB^{dep}$+MA *vs.* MA | ＜0.0001 | 7.793 | 10 |
| Fig. 2a (5-HT) | 5% - 0 Hz *vs.* Ctrl | 0.0659 | 2.065 | 10 |
| Fig. 2a (5-HT) | 10% - 0 Hz *vs.* Ctrl | 0.9653 | 0.04456 | 10 |
| Fig. 2a (5-HT) | 5% - 0.5 Hz *vs.* Ctrl | 0.2237 | 1.297 | 10 |
| Fig. 2a (5-HT) | 10% - 0.5 Hz *vs.* Ctrl | 0.3456 | 0.9897 | 10 |
| Fig. 2a (5-HT) | 5% - 2 Hz *vs.* Ctrl | 0.0890 | 1.883 | 10 |
| Fig. 2a (5-HT) | 10% - 2 Hz *vs.* Ctrl | 0.1118 | 1.744 | 10 |
| Fig. 2a (5-HT) | C48/80 *vs.* Ctrl | ＜0.0001 | 6.639 | 10 |
| Fig. 2a (SP) | 5% - 0 Hz *vs.* Ctrl | 0.7326 | 0.3515 | 10 |
| Fig. 2a (SP) | 10% - 0 Hz *vs.* Ctrl | 0.9298 | 0.09032 | 10 |
| Fig. 2a (SP) | 5% - 0.5 Hz *vs.* Ctrl | 0.4085 | 0.8627 | 10 |
| Fig. 2a (SP) | 10% - 0.5 Hz *vs.* Ctrl | 0.6833 | 0.4201 | 10 |
| Fig. 2a (SP) | 5% - 2 Hz *vs.* Ctrl | 0.7256 | 0.3611 | 10 |
| Fig. 2a (SP) | 10% - 2 Hz *vs.* Ctrl | 0.8463 | 0.1989 | 10 |

| | | | | |
|---|---|---|---|---|
| Fig. 2a (SP) | C48/80 *vs.* Ctrl | ＜0.0001 | 8.476 | 10 |
| Fig. 2b (top) | Ctrl-S *vs.* Ctrl | 0.6125 | 0.5138 | 22 |
| Fig. 2b (top) | Insertion *vs.* Ctrl | 0.4363 | 0.7930 | 22 |
| Fig. 2b (top) | MA *vs.* Insertion | ＜0.0001 | 7.640 | 22 |
| Fig. 2b (top) | CLG+MA *vs*. CLG | 0.1242 | 1.599 | 22 |
| Fig. 2b (top) | CLG+MA *vs*. MA | ＜0.0001 | 6.421 | 22 |
| Fig. 2b (down) | Ctrl-S *vs*. Ctrl | 0.5105 | 0.6690 | 22 |
| Fig. 2b (down) | Insertion *vs*. Ctrl | 0.2427 | 1.200 | 22 |
| Fig. 2b (down) | MA *vs*. Insertion | 0.0017 | 3.564 | 22 |
| Fig. 2b (down) | CLG+MA *vs*. CLG | 0.0525 | 2.050 | 22 |
| Fig. 2b (down) | CLG+MA *vs*. MA | 0.0003 | 4.357 | 22 |
| Fig. 2c | PLL *vs*. Ctrl | 0.6814 | 0.4227 | 10 |
| Fig. 2c | Collagen *vs*. PLL | 0.4556 | 0.7763 | 10 |
| Fig. 2c | Fibronetin *vs*. PLL | ＜0.0001 | 9.776 | 10 |
| Fig. 2d (left) | MA *vs*. Ctrl | ＜0.0001 | 5.341 | 22 |
| Fig. 2d (left) | $FB^{dep}$+MA *vs*. $FB^{dep}$ | 0.3336 | 0.9887 | 22 |
| Fig. 2d (left) | $FB^{dep}$+MA *vs*. MA | ＜0.0001 | 6.457 | 22 |
| Fig. 2d (right) | MA *vs*. Ctrl | 0.0014 | 3.653 | 22 |
| Fig. 2d (right) | $FB^{dep}$+MA *vs*. $FB^{dep}$ | 0.0352 | 2.244 | 22 |
| Fig. 2d (right) | $FB^{dep}$+MA *vs*. MA | 0.0024 | 3.435 | 22 |
| Fig. 3a (left) | 0 Hz *vs*. Ctrl | 0.0556 | 2.165 | 10 |
| Fig. 3a (left) | 0.5 Hz *vs*. Ctrl | 0.0088 | 3.246 | 10 |
| Fig. 3a (left) | 2 Hz *vs*. Ctrl | 0.0183 | 2.815 | 10 |
| Fig. 3a (right) | 0 Hz *vs*. Ctrl | 0.0298 | 2.531 | 10 |
| Fig. 3a (right) | 0.5 Hz *vs*. Ctrl | 0.0040 | 3.722 | 10 |
| Fig. 3a (right) | 2 Hz *vs*. Ctrl | 0.0019 | 4.163 | 10 |
| Fig. 3b (left) | 0 Hz *vs*. Ctrl | 0.0065 | 3.425 | 10 |
| Fig. 3b (left) | 0.5 Hz *vs*. Ctrl | 0.0282 | 2.563 | 10 |

| | | | | |
|---|---|---|---|---|
| Fig. 3b (left) | 2 Hz *vs.* Ctrl | 0.0041 | 3.705 | 10 |
| Fig. 3b (right) | 0 Hz *vs.* Ctrl | 0.0082 | 3.289 | 10 |
| Fig. 3b (right) | 0.5 Hz *vs.* Ctrl | 0.0334 | 2.464 | 10 |
| Fig. 3b (right) | 2 Hz *vs.* Ctrl | 0.0033 | 3.834 | 10 |
| Fig. 3c | No stretch *vs.* Ctrl (8h) | 0.0003 | 5.468 | 10 |
| Fig. 3c | Stretch *vs.* No stretch (8h) | 0.0330 | 2.473 | 10 |
| Fig. 3c | No stretch *vs.* Ctrl (12h) | ＜0.0001 | 80586 | 10 |
| Fig. 3c | Stretch *vs.* No stretch (12h) | 0.0078 | 3.319 | 10 |
| Fig. 3c | No stretch *vs.* Ctrl (16h) | ＜0.0001 | 8.808 | 10 |
| Fig. 3c | Stretch *vs.* No stretch (16h) | 0.0042 | 3.684 | 10 |
| Fig. 3d | Stretch *vs.* Ctrl (NGF) | 0.1193 | 1.704 | 10 |
| Fig. 3d | Stretch *vs.* Ctrl (M-CSF) | 0.3161 | 1.055 | 10 |
| Fig. 3d | Stretch *vs.* Ctrl (CXCL1) | 0.6030 | 0.5371 | 10 |
| Fig. 3d | Stretch *vs.* Ctrl (IL-33) | 0.0020 | 4.148 | 10 |
| Fig. 3d | Stretch *vs.* Ctrl (SCF) | 0.0239 | 2.660 | 10 |
| Fig. 3f (top) | Stretch *vs.* Ctrl | ＜0.0001 | 11.25 | 991 |
| Fig. 3f (top) | Y-27632+stretch *vs.* Stretch | ＜0.0001 | 12.32 | 937 |
| Fig. 3f (down) | Stretch *vs.* Ctrl | ＜0.0001 | 13.02 | 611 |
| Fig. 3f (down) | Y-27632+stretch *vs.* Stretch | ＜0.0001 | 27.15 | 760 |
| Fig. 4a (IL-33 *vs.* 5-HT) | 10 ng/mL *vs.* 0 | 0.5430 | 0.6297 | 10 |
| Fig. 4a (IL-33 *vs.* 5-HT) | 20 ng/mL *vs.* 0 | 0.0006 | 4.910 | 10 |
| Fig. 4a (IL-33 *vs.* 5-HT) | 40 ng/mL *vs.* 0 | 0.0054 | 3.533 | 10 |
| Fig. 4a (IL-33 *vs.* 5-HT) | 50 ng/mL *vs.* 0 | 0.0272 | 2.585 | 10 |
| Fig. 4a (IL-33 *vs.* 5-HT) | 60 ng/mL *vs.* 0 | ＜0.0001 | 6.355 | 10 |
| Fig. 4a (IL-33 *vs.* 5-HT) | 100 ng/mL *vs.* 0 | 0.0853 | 1.909 | 10 |
| Fig. 4a (IL-33 *vs.* SP) | 10 ng/mL *vs.* 0 | 0.3128 | 1.063 | 10 |
| Fig. 4a (IL-33 *vs.* SP) | 20 ng/mL *vs.* 0 | 0.0044 | 3.663 | 10 |
| Fig. 4a (IL-33 *vs.* SP) | 40 ng/mL *vs.* 0 | 0.0011 | 4.525 | 10 |

| | | | | |
|---|---|---|---|---|
| Fig. 4a (IL-33 *vs.* SP) | 50 ng/mL *vs.* 0 | 0.0069 | 3.388 | 10 |
| Fig. 4a (IL-33 *vs.* SP) | 60 ng/mL *vs.* 0 | 0.0010 | 4.559 | 10 |
| Fig. 4a (IL-33 *vs.* SP) | 100 ng/mL *vs.* 0 | 0.9875 | 0.01608 | 10 |
| Fig. 4a (SCF *vs.* 5-HT) | 20 ng/mL *vs.* 0 | 0.0038 | 3.749 | 10 |
| Fig. 4a (SCF *vs.* 5-HT) | 40 ng/mL *vs.* 0 | 0.0106 | 3.135 | 10 |
| Fig. 4a (SCF *vs.* 5-HT) | 50 ng/mL *vs.* 0 | 0.4207 | 0.8396 | 10 |
| Fig. 4a (SCF *vs.* 5-HT) | 60 ng/mL *vs.* 0 | 0.2771 | 1.149 | 10 |
| Fig. 4a (SCF *vs.* 5-HT) | 100 ng/mL *vs.* 0 | 0.2358 | 1.261 | 10 |
| Fig. 4a (SCF *vs.* SP) | 20 ng/mL *vs.* 0 | 0.3556 | 0.9685 | 10 |
| Fig. 4a (SCF *vs.* SP) | 40 ng/mL *vs.* 0 | 0.5936 | 0.5512 | 10 |
| Fig. 4a (SCF *vs.* SP) | 50 ng/mL *vs.* 0 | 0.1239 | 1.680 | 10 |
| Fig. 4a (SCF *vs.* SP) | 60 ng/mL *vs.* 0 | 0.0222 | 2.702 | 10 |
| Fig. 4a (SCF *vs.* SP) | 100 ng/mL *vs.* 0 | 0.0156 | 2.907 | 10 |
| Fig. 4b (left) | MA *vs.* Ctrl-S | ＜0.0001 | 5.112 | 22 |
| Fig. 4b (left) | ISCK-5 *vs.* Ctrl-S | 0.0142 | 2.665 | 22 |
| Fig. 4b (left) | ISCK-10 *vs.* Ctrl-S | 0.0092 | 2.857 | 22 |
| Fig. 4b (left) | ISCK-5+MA *vs.* MA | ＜0.0001 | 5.606 | 22 |
| Fig. 4b (left) | ISCK-10+MA *vs.* MA | ＜0.0001 | 6.753 | 22 |
| Fig. 4b (left) | SCF *vs.* Ctrl-T | ＜0.0001 | 5.877 | 22 |
| Fig. 4b (right) | MA *vs.* Ctrl-S | ＜0.0001 | 7.227 | 22 |
| Fig. 4b (right) | ISCK-5 *vs.* Ctrl-S | 0.0906 | 1.770 | 22 |
| Fig. 4b (right) | ISCK-10 *vs.* Ctrl-S | 0.0061 | 3.034 | 22 |
| Fig. 4b (right) | ISCK-5+MA *vs.* MA | ＜0.0001 | 15.91 | 22 |
| Fig. 4b (right) | ISCK-10+MA *vs.* MA | ＜0.0001 | 12.79 | 22 |
| Fig. 4b (right) | SCF *vs.* Ctrl-T | 0.1849 | 1.369 | 22 |
| Fig. 4c (left) | 0.1 μg/mL *vs.* 0 | 0.0007 | 4.788 | 10 |
| Fig. 4c (left) | 0.5 μg/mL *vs.* 0 | 0.0195 | 2.779 | 10 |
| Fig. 4c (left) | 1.0 μg/mL *vs.* 0 | 0.0273 | 2.583 | 10 |

| | | | | |
|---|---|---|---|---|
| Fig. 4c (right) | 0.1 μg/mL *vs.* 0 | 0.9314 | 0.8833 | 10 |
| Fig. 4c (right) | 0.5 μg/mL *vs.* 0 | 0.0006 | 4.978 | 10 |
| Fig. 4c (right) | 1.0 μg/mL *vs.* 0 | 0.0015 | 4.313 | 10 |
| Fig. 4d (left) | MA-T *vs.* Ctrl-T | 0.0006 | 4.032 | 22 |
| Fig. 4d (left) | IL-33 *vs.* Ctrl-T | 0.0029 | 3.348 | 22 |
| Fig. 4d (left) | MA-S *vs.* Ctrl-S | 0.0014 | 3.653 | 22 |
| Fig. 4d (left) | sST2+MA *vs.* MA-S | 0.0003 | 4.293 | 22 |
| Fig. 4d (right) | MA-T *vs.* Ctrl-T | ＜0.0001 | 6.799 | 22 |
| Fig. 4d (right) | IL-33 *vs.* Ctrl-T | ＜0.0001 | 7.365 | 22 |
| Fig. 4d (right) | MA-S *vs.* Ctrl-S | ＜0.0001 | 5.577 | 22 |
| Fig. 4d (right) | sST2+MA *vs.* MA-S | ＜0.0001 | 8.586 | 22 |
| Fig. 4e | MA-T *vs.* Ctrl-T | 0.0012 | 4.448 | 10 |
| Fig. 4e | IL-33 *vs.* Ctrl-T | 0.0001 | 6.020 | 10 |
| Fig. 4e | MA-S *vs.* Ctrl-S | ＜0.0001 | 9.830 | 10 |
| Fig. 4e | sST2+MA *vs.* MA-S | ＜0.0001 | 7.050 | 10 |
| Fig. 5b (IL-1β) | MA-T *vs.* Ctrl-T | ＜0.0001 | 7.801 | 10 |
| Fig. 5b (IL-1β) | IL-33 *vs.* Ctrl-T | 0.0007 | 4.856 | 10 |
| Fig. 5b (IL-1β) | SCF *vs.* Ctrl-T | 0.4897 | 0.7171 | 10 |
| Fig. 5b (IL-6) | MA-T *vs.* Ctrl-T | 0.0013 | 4.408 | 10 |
| Fig. 5b (IL-6) | IL-33 *vs.* Ctrl-T | 0.0060 | 3.473 | 10 |
| Fig. 5b (IL-6) | SCF *vs.* Ctrl-T | 0.6673 | 0.4428 | 10 |
| Fig. 5b (TNF-α) | MA-T *vs.* Ctrl-T | 0.0117 | 3.078 | 10 |
| Fig. 5b (TNF-α) | IL-33 *vs.* Ctrl-T | 0.0321 | 2.489 | 10 |
| Fig. 5b (TNF-α) | SCF *vs.* Ctrl-T | 0.5241 | 0.6602 | 10 |
| Fig. 5d (IL-1β) | MA-S *vs.* Ctrl-S | ＜0.0001 | 7.313 | 10 |
| Fig. 5d (IL-1β) | sST2+MA *vs.* MA-S | 0.0112 | 3.102 | 10 |
| Fig. 5d (IL-6) | MA-S *vs.* Ctrl-S | ＜0.0001 | 18.46 | 10 |
| Fig. 5d (IL-6) | sST2+MA *vs.* MA-S | 0.0040 | 3.722 | 10 |

| | | | | |
|---|---|---|---|---|
| Fig. 5d (TNF-α) | MA-S *vs.* Ctrl-S | 0.0015 | 4.319 | 10 |
| Fig. 5d (TNF-α) | sST2+MA *vs.* MA-S | 0.0028 | 3.944 | 10 |
| Extended Data Fig. 7c (left) | Stretch *vs.* Ctrl | 0.0072 | 5.056 | 4 |
| Extended Data Fig. 7 (right) | Stretch *vs.* Ctrl | 0.0016 | 7.666 | 4 |
| Extended Data Fig. 8a | 10 ng/mL *vs*. 0 | 0.0053 | 3.552 | 10 |
| Extended Data Fig. 8a | 20 ng/mL *vs*. 0 | 0.0102 | 3.159 | 10 |
| Extended Data Fig. 8a | 50 ng/mL *vs*. 0 | 0.0705 | 2.024 | 10 |
| Extended Data Fig. 8b | 1 μM *vs*. 0 | 0.0004 | 5.241 | 10 |
| Extended Data Fig. 8b | 5 μM *vs*. 0 | ＜0.0001 | 7.971 | 10 |
| Extended Data Fig. 8b | 10 μM *vs*. 0 | ＜0.0001 | 9.423 | 10 |
| Extended Data Fig. 8b | 20 μM *vs*. 0 | 0.0073 | 3.359 | 10 |
